# Widespread hybridization between invasive bleak (*Alburnus alburnus*) and native Iberian chubs (*Squalius* spp.): a neglected threat

**DOI:** 10.1101/2022.03.04.483063

**Authors:** M Morgado-Santos, M Curto, CM Alexandre, MJ Alves, HF Gante, C Gkenas, JP Medeiros, PJ Pinheiro, PR Almeida, MF Magalhães, F Ribeiro

## Abstract

Hybridization between native and exotic species is a major conservation issue. In Iberian rivers, which are simultaneously among the most invaded and diverse ecosystems, harboring a high proportion of endemic fishes, this issue is becoming highly concerning. To date, hybridization has been reported to occur between the invasive bleak *Alburnus alburnus* and the endemic chubs *Squalius alburnoides* and *S. pyrenaicus*, in some scattered locations. However, the bleak is increasingly spreading in the region, potentially increasing the risks of hybridization with these and other *Squalius* species. To gather a more comprehensive picture on the current extension of hybridization, we compiled and mapped records on hybrids between bleak and chubs collected by multiple research teams in different ongoing projects in the Portuguese territory, and genetically assessed the hybrid identity of specimens in an area of sympatry between bleak and *S. carolitertii*, using nuclear and mitochondrial markers. We found that hybridization with bleak is widespread in several Portuguese river basins and involves at least three endemic chubs, *S. alburnoides, S. pyrenaicus* and *S. carolitertii*. This may have serious consequences, including not only waste of reproductive effort and damage of genetic integrity, but also putative shifts on the reproductive dynamics of the *Squalius* system, which includes endemic hybrids, reproducing sexually and nonsexually. Hybridization should be recognized as a significant impact of biological invasions, and we recommend that future studies should characterize the fitness of hybrids and their ecological and genetic interactions with parental Iberian fish, to elucidate effective conservation measures.

## INTRODUCTION

Biological invasions threaten biodiversity and contribute to species loss through different mechanisms, including hybridization (Lodge 1993, Olden et al. 2004, Vellend et al. 2007, Bellard et al. 2016). Interbreeding between exotic species colonizing new areas and local native species increases extinction risk of the latter, with conservation concerns even if hybrids are unviable or infertile due to waste of reproduction effort (Rhymer & Simberloff 1996, Allendorf et al. 2001, Largiadèr 2007, Ribeiro & Leunda 2012). Fertile hybrids may form homoploid or polyploid lineages, potentially displacing parental species and leading to admixture between native and exotic genomes (Dowling & Secor 1997, Perry et al. 2001, Largiadèr 2007, Quilodrán et al. 2015). The introgression of non-native genes into native species is irreversible, damaging genetic integrity and disrupting local adaptations through the introduction of maladaptive genes, possibly causing the extinction of native genotypes (Rhymer & Simberloff 1996, Huxel 1999, Mack et al. 2000, Fitzpatrick et al. 2010). Conversely, exotic species may benefit from introgression of genetic variability and locally adapted genes from the native species, alleviating the loss of genetic variation through founder effect (Rhymer & Simberloff 1996, Huxel 1999, Currat et al. 2008, Hall & Ayres 2009).

Hybridization between native and exotic species is particularly concerning in Iberian streams, which are among the most biodiverse and invaded ecosystems in the world (Leprieur et al. 2008). In particular, fishes are highly prone to interbreeding due to external fertilization, weak reproductive isolation, high interspecific genetic compatibility and human-mediated decrease in habitat complexity (Verspoor & Hammar 1991, Scribner et al. 2000, Docker et al. 2003, Chávez & Turgeon 2007). The spread of non-native alleles through introgression among the gene pools of Iberian fishes may have serious consequences on the fitness, ecology, behavior and likelihood of population persistence (Rhymer & Simberloff 1996, Huxel 1999, Ribeiro & Leunda 2012).

First introduced to Iberian streams in 1992, the bleak *Alburnus alburnus* has been expanding since, being currently widespread and locally abundant in most Portuguese river basins (Latorre et al. 2018, Collares-Pereira et al. 2021, Martelo et al. 2021). Hybridization between *A. alburnus* and Iberian chubs *Squalius pyrenaicus* and the allopolyploid complex *Squalius alburnoides* has been reported in a few localities (Almodóvar et al. 2012, Sousa-Santos et al. 2018), but the extent of interbreeding with these and potentially other *Squalius* species, such as *Squalius carolitertii*, remains unknown. Clarification is particularly important, as the endemic *Squalius* system naturally includes hybrids that reproduce sexually and nonsexually (i.e., *S. alburnoides*), and their reproductive dynamics may be disturbed by the inclusion of another hybridizing species (i.e., the exotic *A. alburnus*), compromising the survival of the native populations. Here, we sought to assess interbreeding between the invasive bleak and Iberian chubs across Portuguese river basins. We mapped the occurrence of hybrids identified using morphological characters, building on records gathered from multiple research teams. Hybridization between bleak and *Squalius carolitertii* was further analyzed by molecular analysis of individuals with intermediate morphology, captured in areas of sympatry of the two species. Results provided a more comprehensive picture regarding the distribution of hybridization risks associated with bleak invasion throughout the Portuguese river basins, and were used to open new perspectives on the consequences of hybridization associated with biological invasions.

## MATERIALS AND METHODS

### Occurrence data and mapping

Data on the occurrence of hybrids between *A. alburnus* and *Squalius* spp. were collected between 2015 and 2021 throughout the distribution range of *A. alburnus*, covering areas of sympatry with *S. alburnoides, S. pyrenaicus* and *S. carolitertii* in Portugal. Records were obtained in the frame of several research projects (see acknowledgments), generally from electrofishing (300 V, 1-2 A). Hybrids were identified using morphological traits, focusing on meristic characters recommended by Almodóvar et al. (2012), namely number of: a) lateral line scales; b) transverse scales; c) branched dorsal fin rays; d) branched ventral fin rays; and e) branched anal fin rays. Geographic locations of hybrid records were mapped using Google Maps Pro (v. 7.3.4.8248).

### Fish sampling and molecular analysis of hybrids

Additional fish sampling was performed in August 2020 and in June 2021, using electrofishing (300 V, 1-2 A), at two sites in Vizela River (Ave basin; 41.375300, - 8.333808 and 41.375189, -8.264944), where both *A. alburnus* and *S. carolitertii* are abundant. Five putative hybrids (morphologically identified as described above) were fin clipped for further molecular analysis. Invasive *A. alburnus* and hybrids were euthanized with a lethal dose of anesthetic (MS-222), and stored in 4% formaldehyde for later integration in the collections of the National Museum of Natural History and Science, University of Lisbon. Native fish were returned to the river.

Total DNA was extracted from the fin clips of the hybrids with a commercial isolation kit (E.Z.N.A. Tissue DNA Kit, Omega Bio-Tek, USA), following manufacturer’s instructions, checked for purity and concentration (ng/uL) in Nanodrop (Thermo Fisher Scientific), and stored at –20ºC. One nuclear and one mitochondrial loci (beta-actin and COI, respectively) were amplified and sequenced for each putative hybrid individual, to assess genomic composition and maternal inheritance.

Beta-actin was amplified using the primers described in Sousa-Santos et al. (2005), and COI was amplified with a universal mix of four different primers, namely VF2_t1, FishF2_t1, FishR2_t1 and FR1d_t1, with M13 tails to facilitate sequencing, as described in Ivanova et al. (2007). PCRs were performed with final concentrations of 5– 15 ng/uL of DNA template, 2 mM of MgCl_2_, 0.1–0.2 mM of each dNTP and 0.03–0.05 U/μL of DNA polymerase, namely *GoTaq* (Promega, USA) or *Taq PCR Mix with MgCl2* (Abnova, Taiwan). PCR conditions for both genes were as follows: 1 cycle of 95ºC for 5 min (initial denaturation), 35 cycles of 95ºC for 30 s (denaturation), 55ºC for 40 s (annealing) and 72ºC for 90 s (elongation), and 1 cycle of 72ºC for 10 min (final elongation). A sample of each PCR product was run in 3% agarose gels to check for successful amplification, using EZ-Vision Bluelight DNA dye (VWR Life Sciences, USA), purified using ExoCleanUp FAST PCR clean-up reagent (VWR Life Sciences USA), sequenced at the forward direction in outsourcing at StabVida (Portugal) and deposited in GenBank.

Sequences of beta-actin and COI from the parental species (*A. alburnus* and *S. carolitertii*) were obtained from GenBank, aligned in BioEdit and used to create consensus sequences, to compare SNPs between the putative hybrids and the parental species. The alleles of the beta-actin sequences of heterozygous individuals were extracted using Indigo (Gear Genomics), and COI sequences were blasted in BOLD Systems (The Barcode of Life Data System). The GenBank accession numbers of the sequences used are listed in the Supplementary Table 1.

### Ethical statement

All field and laboratorial procedures followed the recommended ethical guidelines and legislation regarding animal capture, manipulation and experimentation for scientific purposes, and were conducted under permits obtained from the Portuguese Nature Conservation Authority (ICNF – Instituto da Conservação da Natureza e das Florestas).

## RESULTS

Hybrids were found across seven river basins, in areas of sympatry between *A. alburnus* and *S. alburnoides, S. pyrenaicus*, and *S. carolitertii*, namely in the Ave, Douro, Vouga, Mondego, Tagus, Sado and Guadiana river basins (Fig. 1).

**Figure 1.**
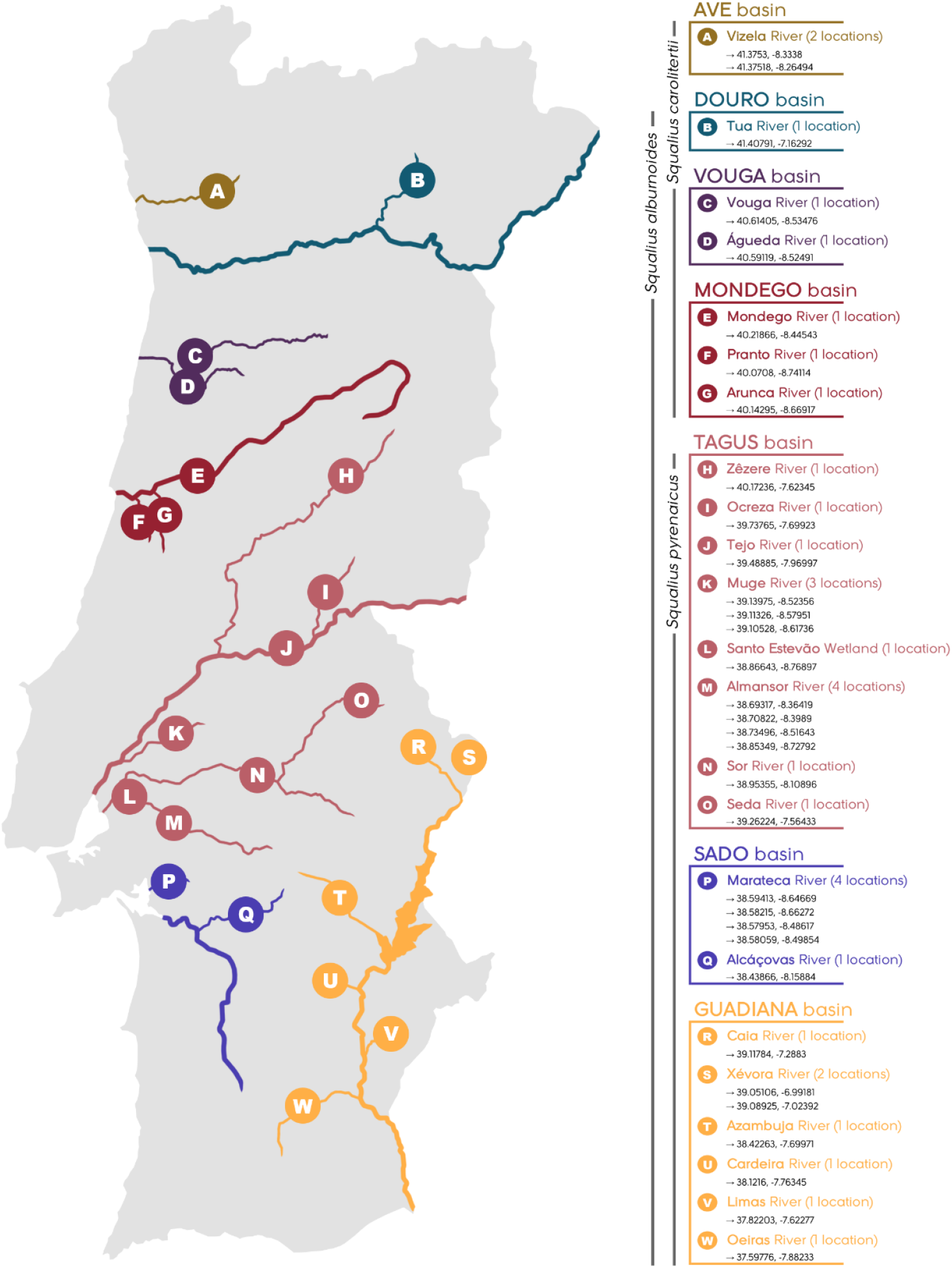
Map of hybrid records in Portuguese river basins. Geographical coordinates [World Geodetic System (WGS84), in decimal degrees] of capture localities are shown for each basin and river. The areas of sympatry between *A. alburnus* and each *Squalius* species where hybrids have been found are represented by the gray vertical lines on the left of the list of basins/rivers.

Sequencing of the beta-actin and COI genes confirmed the hybrid identity of the five putative hybrids between *A. alburnus* and *S. carolitertii* collected in the Ave river basin (Vizela River) (Table I; see sequences’ chromatograms in the Supplementary Material). All were mothered by *S. carolitertii* and fathered by *A. alburnus*, as indicated through direct analyses of diagnostic SNPs, and by blasting COI sequences in BOLD system databases.

**Table I.**
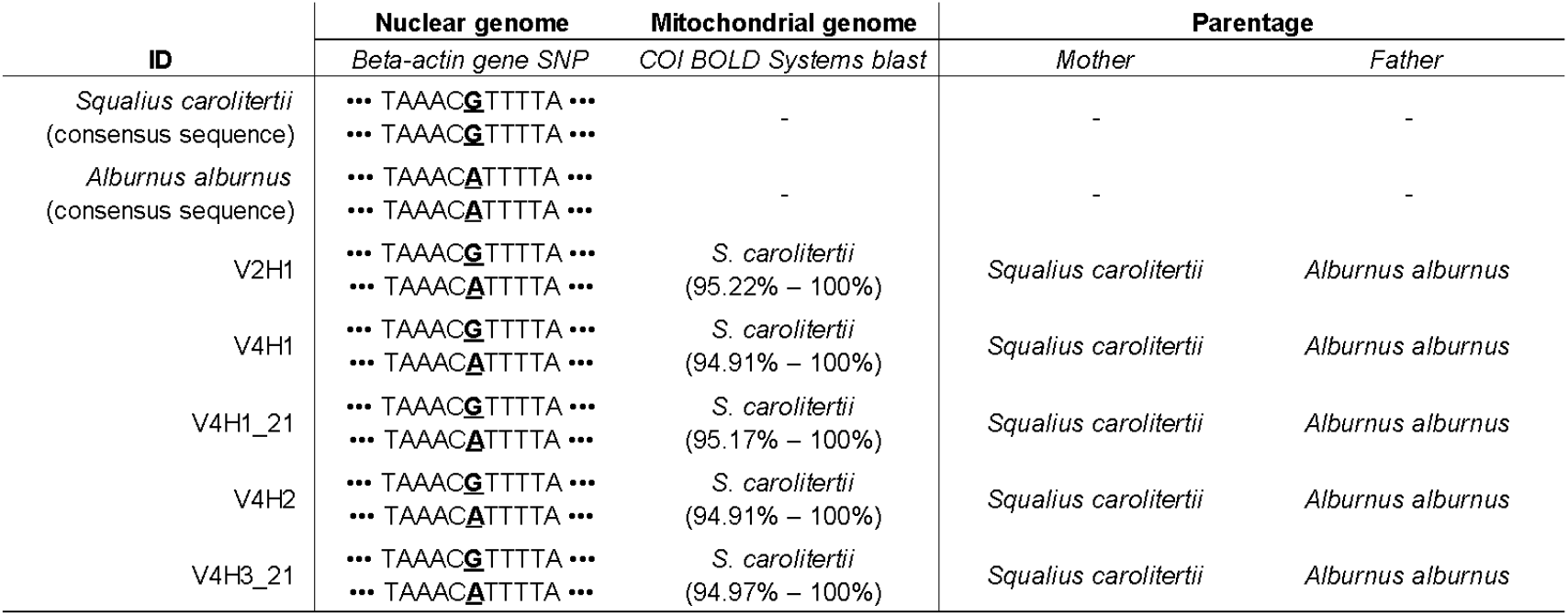
Results of SNP analyses of beta-actin nuclear gene and of BOLD blast of COI mitochondrial gene, discriminating the parental species involved and the direction of hybridization for each hybrid analyzed (ID codes V2H1, V4H1, V4H1_21, V4H2, V4H3_21). Low and top percentages of similarity in BOLD blasts are shown under each COI identification.

## DISCUSSION

Results from the current study show that hybridization between *A. alburnus* and endemic *Squalius* occurs in areas of sympatry, in seven major river basins across Portugal, indicating that hybridization is likely widespread in the region. This is consistent with previous evidence that hybridization may become widespread, with hybrids occurring frequently or even massively, when interbreeding between exotic and native species leads to viable hybrid offspring (Huxel 1999, Rubidge et al. 2001, Meldgaard et al. 2007, Meraner et al. 2013, Buonerba et al. 2015).

Hybridization between *A. alburnus* and *Squalius* spp. appears to be more extensive than previously thought. In addition to interbreeding with *S. alburnoides* and *S. pyrenaicus*, as previously reported for the Sado and Guadiana basins (Sousa-Santos et al. 2018) and elsewhere (Almodóvar et al. 2012), *A. alburnus* also interbreeds and can produce viable hybrids with *S. carolitertii* in northern basins (e.g., Mondego, Vouga, Douro and Ave). Incomplete reproductive isolation between the invasive bleak and Iberian chubs raises significant conservation concerns. Indeed, as *A. alburnus* shows protracted spawning (Latorre et al. 2018) and increasingly spreads across the Iberian Peninsula, it cannot be ignored that it may interbreed with critically endangered chubs with very restricted distributions, which are likely to be severely impacted by hybridization, namely *S. aradensis, S. castellanus, S. laietanus, S. malacitanus, S. palaciosi, S. torgalensis* and *S. valentinus* (Doadrio 2001, Rogado et al. 2005).

Although the fertility of the hybrids remains unknown, hybridization with *A. alburnus* is likely to cause impacts on native chubs even if hybrids are sterile, due to the waste of reproductive effort. Although *Squalius* spp. are multiple spawners, the number of egg batches laid per reproductive season is limited. For instance, *S. alburnoides* females lay between 1 and 6 egg batches per year (Morgado-Santos et al. 2016), and *S. pyrenaicus* and *S. carolitertii* have more restricted spawning seasons (Pires et al. 2000; Alexandre et al. 2018). Hybridization with the invasive bleak will be particularly concerning if it is mainly unidirectional between *S. carolitertii* females and *A. alburnus* males, as suggested in our results, considering *S. pyrenaicus* and *S. carolitertii* bisexual species are already sexually parasitized by the allopolyploid *S. alburnoides* (Collares-Pereira et al. 2013).

The *S. alburnoides* allopolyploid complex includes several forms with different ploidy combinations of the parental species’ genomes (genomotypes), involved in intricate reproductive dynamics. The interbreeding with the invasive bleak may disrupt this dynamics, with important conservation consequences. *Squalius alburnoides* genomotypes are interdependent, with the production of each genomotype relying on crosses involving other genomotypes, including the parental bisexual species (Collares-Pereira et al. 2013). The intricate reproductive network of the *Squalius* system is uphold by crosses with variable frequencies occurring among specific genomotypes, which may lead populations to an equilibrium and evolutionary stability, maintaining them at a stable hybrid state, or rout them towards hybrid speciation (Morgado-Santos et al. 2015). The inclusion of the interbreeding invasive bleak into the *Squalius* system may change this reproductive network, potentially causing imbalances in the genomotype composition of populations, which may ultimately increase extinction risk. Shifts in the reproductive dynamics of native hybrids due to the introduction of an exotic species has been already reported for the *Pelophylax* hybrid system (Quilodrán et al. 2015), which shares many traits with the *Squalius* system.

Hybridization between species of the *Squalius* system and *A. alburnus* may also lead to genetic deterioration of the native genomes, which may be particularly detrimental for *S. alburnoides*. This allopolyploid fish includes a genome of an extinct species, that currently is expressed without recombining with *Squalius* genomes (Collares-Pereira et al 2013). However, the genome of the exotic *A. alburnus* is phylogenetically close to this extinct genome (Perea et al. 2010), and thus meiotic recombination may be more likely to occur. While fitness benefits cannot be excluded at present, admixture could more likely disrupt local adaptations, lead to the introduction of maladaptive genes, and strongly affect the reproductive dynamics of the allopolyploid complex, because mate choice by *S. alburnoides* females is influenced by the genetic integrity of their own genomes and of those of the available mates (Morgado-Santos et al. 2018).

Finally, we highlight the importance of early detection of hybrids, especially for species already threatened, because introgression may occur quickly and even be favored by natural selection (Echelle & Connor 1989, Fitzpatrick et al. 2010, Kovach et al. 2015). Hybrids identified using meristic characters and confirmed by sequencing a few diagnostic loci are more likely to be F1 or hybrids with highly admixed genomes, which have a higher probability of showing obvious intermediate morphology and heterozygosity of diagnostic SNPs. Introgressed individuals via backcrossing may be easily missed using these methods, particularly if mainly focusing on morphological traits (Haynes et al. 2011), thereby underestimating the extent of introgressive hybridization. Such limitations can be overcome by complementing morphological identification with genome-wide molecular approaches, such as RAD-sequencing (Hohenlohe et al. 2011), SNP panels (von Thaden et al. 2020) or SSR-GBS (Curto et al. 2019), tools that are becoming increasingly available.

## CONCLUSION

Biological invasions have deserved increasing attention in the last decades, but studies seldom focus on the inconspicuous and silent consequences of hybridization between native and exotic species. The widespread distribution of hybrids between invasive *A. alburnus* and several *Squalius* species across Portuguese river basins suggests that hybridization is likely to occur frequently, with potentially high impacts on endemic and endangered fishes. Waste of reproductive effort by interbreeding with the invader is likely to accentuate the decline of populations that are already threatened. Interbreeding with bleak may interfere with several aspects of the reproductive network of the *Squalius alburnoides* system, leading to the loss of a stable structure. Finally, if hybrids are fertile and reproduce sexually (with meiotic recombination), native and exotic genomes may admix, irreversibly altering the genetic composition of native species. This aspect needs to be further investigated to evaluate the severity of ongoing hybridization. Given these potentially serious impacts and the continued expansion of *A. alburnus*, we recommend that future studies integrate molecular tools and characterize hybrid fitness, and their ecological and genetic interactions with parental Iberian fish, to elucidate effective conservation measures for Iberian fishes.

## Supporting information

Hybrid sequences chromatograms

## ACKNOWLEDGEMENTS

We thank all members of the research projects from the Foundation for Science and Technology (FCT), which provided data for this study, namely FRISK (PTDC/AAG-MAA/0350/2014), ISO-INVA (PTDC/CTA-AMB/29105/2017), ENVMETAGENOMICS (PTDC/BIA-CBI/31644/2017) and SONICINVADERS (PTDC/CTA-AMB/ 28782/2017). Additionally, Portuguese national funds were received from the Foundation for Science and Technology through the strategic plan of the Marine and Environmental Sciences Centre (MARE) (UID/Multi/04326/2019) and through the individual contract attributed to Carlos M. Alexandre within the project CEECIND/02265/2018. We thank the Fish Invasions Lab team members and students for helping on field work and laboratorial procedures.

## Notes

### Competing Interest Statement

The authors have declared no competing interest.

